# The phycobilisome linker protein ApcG interacts with photosystem II and regulates energy transfer to photosystem I in *Synechocystis sp.* PCC 6803

**DOI:** 10.1101/2023.05.22.541798

**Authors:** Roberto Espinoza-Corral, Masakazu Iwai, Tomáš Zavřel, Sigal Lechno-Yossef, Markus Sutter, Jan Červený, Krishna K. Niyogi, Cheryl A. Kerfeld

**Affiliations:** MSU-DOE Plant Research Laboratory, Michigan State University, East Lansing, MI, USA; Environmental Genomics and Systems Biology Division, Lawrence Berkeley National Laboratory, Berkeley, CA, USA; Molecular Biophysics and Integrated Bioimaging Division, Lawrence Berkeley National Laboratory, Berkeley, CA, USA; Department of Biochemistry and Molecular Biology, Michigan State University, East Lansing, MI, USA; Department of Plant and Microbial Biology, University of California, Berkeley, CA, USA; Howard Hughes Medical Institute, University of California, Berkeley, CA, USA; Department of Adaptive Biotechnologies, Global Change Research Institute of the Czech Academy of Sciences, Brno, Czech Republic

**Keywords:** Phycobilisomes, Photosystem, *Synechocystis*, State transitions, light harvesting

## Abstract

Photosynthetic organisms harvest light using pigment-protein super-complexes. In cyanobacteria, these are water-soluble antennae known as phycobilisomes (PBSs). The light absorbed by PBS is transferred to the photosystems in the thylakoid membrane to drive photosynthesis. The energy transfer between these super-complexes implies that protein-protein interactions allow the association of PBS with the photosystems. However, the specific proteins involved in the interaction of PBS with the photosystems are not fully characterized. Here, we show that the newly discovered PBS linker protein ApcG interacts specifically with photosystem II through its N-terminal region. Growth of cyanobacteria is impaired in *apcG* deletion strains under light-limiting conditions. Furthermore, complementation of these strains using a phospho-mimicking version of ApcG exhibit reduced growth under normal growth conditions. Interestingly, the interaction of ApcG with photosystem II is affected when a phospho-mimicking version of ApcG is used, targeting the positively charged residues interacting with thylakoid membrane suggesting a regulatory role mediated by phosphorylation of ApcG. Low temperature fluorescence measurements showed increased photosystem I fluorescence in *apcG* deletion and complementation strains. The photosystem I fluorescence was the highest in the phospho-mimicking complementation strain while pull-down experiment showed no interaction of ApcG with PSI under any tested condition. Our results highlight the importance of ApcG for selectively directing energy harvested by the PBS and implies that the phosphorylation status of ApcG plays a role in regulating energy transfer from PSII to PSI.

## INTRODUCTION

Light harvesting in cyanobacteria and red algae is enhanced by soluble protein-pigment super-complexes known as phycobilisomes (PBSs). The energy absorbed by PBSs is transferred to photosystem II (PSII) and photosystem I (PSI) embedded in the thylakoid membrane. In the model cyanobacterium *Synechocystis sp.* PCC 6803 (hereafter referred to as *Synechocystis*), the PBS consists of a tri-cylindrical core with six attached rods, in a hemidiscoidal arrangement (Dominguez-Martin et al., 2022, Gantt and Conti, 1969, Bryant et al., 1979). The PBS consist of phycobiliproteins containing covalently linked bilin pigments and assembled into disc-like trimers (αβ)_3_ or hexamers (αβ)_6_, and colorless linker proteins that connect the phycobiliprotein discs (Tandeaudemarsac and Cohenbazire, 1977, Adir, 2005, Glauser et al., 1992, Anderson and Toole, 1998). Their size and the ability to absorb light at wavelengths between 550 to 650 nm, where chlorophyll *a* absorption is low, increases the area and the spectral range of light harvesting in cyanobacteria.

The energy transfer from water-soluble PBS to the photosystems entails a close contact between these super-complexes. A recent structure of the PBS-PSII super-complex (with a resolution of 14.3 Å) from the red algae *Porphyridium purpureum* UTEX 2757 provides insights on the architecture of this interaction, involving allophycocyanin E (ApcE, also classified as core-membrane linker protein L_CM_), allophycocyanin D (ApcD) and three unidentified connector proteins from PBS (Li et al., 2021). These phycobiliproteins have been classified as terminal emitters responsible for energy transfer to PSII (ApcE) and PSI (ApcD) (Gindt et al., 1992, Dong et al., 2009, Peng et al., 2014, Liu et al., 2013). Additionally, pull-down experiments followed by mass-spectrometry of PBS and photosystems from *Synechocystis* have shown putative contact sites in the interface of these super-complexes involving also ApcD and ApcE (Liu et al., 2013).

Furthermore, PBSs and photosystems have been observed to be organized into three main microdomains in the thylakoid membrane, one containing the trimeric PSI, another one with the dimeric PSII and PBS, and the last one including both photosystems and PBS (Straskova et al., 2019). Additionally, cryogenic electron tomography of *Synechocystis* has shown that PBSs organize in arrays on the thylakoid membrane, presumably increasing the efficiency of light harvesting (Rast et al., 2019). While PBSs are able to transfer energy to either PSII or PSI, the proteins involved in PBS-PSII or PBS-PSI interactions that govern such specificity are still unknown. Moreover, energy transfer from PBS to PSI could follow two plausible models. On the one hand there is the model of PBS mobility on thylakoid membranes that assumes detachment of PBS from PSII to then attach to PSI (Sarcina et al., 2001, Mullineaux et al., 1997, Yang et al., 2007). On the other hand, the spillover model proposes that the energy absorbed by PBS is first transferred to PSII, which then transfers the excess of energy to PSI (McConnell et al., 2002, Folea et al., 2008, Olive et al., 1997). There is an efficient mechanism for short timescales regulation of photosystems activity in response to variations in both the quality and quantity of light known as state transitions. This process aims to prevent photodamage caused by the saturation of the photosynthetic electron transport chain, ensuring a balanced activity of both PSI and PSII. In contrast to plants, there is little consensus on how state transitions are achieved and regulated in cyanobacteria (Calzadilla and Kirilovsky, 2020).

The *Synechocystis* PBS structure recently obtained by cryogenic electron microscopy (Cryo-EM, with a resolution of 2.1-3.5 Å) included different rod conformations and revealed a novel PBS linker protein (ApcG) located at the bottom two cylinders of the tri-cylindrical core (Dominguez-Martin et al., 2022). Due to its location in the PBS core, it has been hypothesized that this linker protein could interact with the photosystems allowing the tethering of PBS to the thylakoid membrane.

Here we show that the PBS linker protein ApcG binds specifically to PSII via its N-terminal region and that this interaction is likely affected by the phosphorylation state of ApcG. Furthermore, an *apcG* deletion strain shows slower growth compared to wild type under light-limiting conditions. Under normal light conditions only a phospho-mimicking *apcG* complementation strain shows slower growth. This phenotype was further characterized by low temperature fluorescence, revealing an imbalance in PSII and PSI activity in the deletion and complementation strains, which exhibited higher PSI fluorescence with the highest in the phospho-mimicking version of ApcG. Our results indicate that ApcG plays a crucial role in PBS-PSII interaction and the transfer of energy towards PSI, presumably via the “spillover” mechanism.

## RESULTS

### Domain organization of ApcG

The recent *Synechocystis* PBS structure showed that two ApcG molecules bind to the PBS at the membrane-facing side, one at each of the bottom core cylinders via the C-terminal domain of ApcG (**Figure 1A, B**) (Dominguez-Martin et al., 2022). The PBS-binding domain is characterized by a conserved FxxM motif, which interdigitates with ApcA at the bottom core cylinder (Liu, 2023) (**Figure 1A, C**). The location observed in the Cryo-EM structure indicates that the N-terminal portion of ApcG extends outwards from the PBS core, presumably to interact with the photosystems in the thylakoid membrane. A sequence search indicated that ApcG homologues are found in 94% of cyanobacterial genomes that also have an ApcE orthologue (Dominguez-Martin et al., 2022). The amino acid conservation of ApcG orthologs from various cyanobacteria species show three conserved domains; i) N-terminal domain, ii) a positively charged middle domain and iii) the PBS binding domain (**Figure 1C**). Additionally, phospho-proteomic data show that ApcG contains several phosphorylation sites (Angeleri et al., 2016). They are located close to the positively charged middle domain, which could play a role in regulating this region’s interaction with the thylakoid membrane (**Figure 1B, D**). Specifically, residues 46-48 (TTS) were found to be phosphorylated under low and high carbon conditions (Angeleri et al., 2016). Furthermore, AlphaFold2 (Jumper et al., 2021) structure prediction of ApcG shows that the middle domain is unstructured compared to the PBS-binding domain (**Figure 1D**). The conservation of ∼20 residues at the N-terminus (**Figure 1C**) suggests that it could play a role in regulating the interaction of PBS with one of the photosystems.

**Figure 1.**
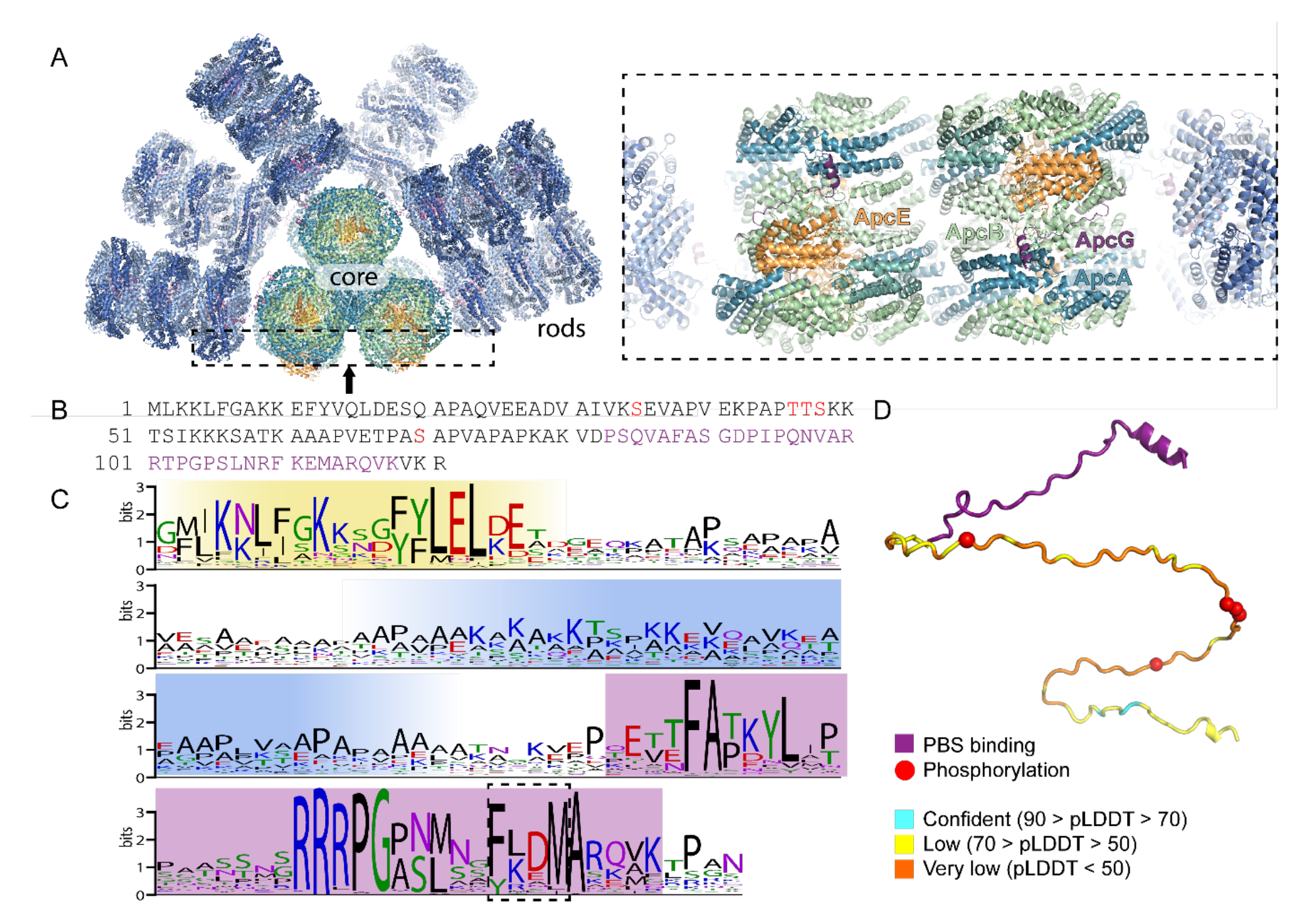
PBS linker protein ApcG binding site, conservation and domain structure. **(A)** Overview of *Synechocystis* PBS (left) and zoomed in view of ApcG binding site at the bottom cylinder (right). **(B)** Primary structure of *Synechocystis* ApcG. Phosphorylation sites are highlighted in red while PBS binding region is in purple. **(C)** Sequence conservation logo of 347 cyanobacterial ApcG homologs. The N-terminal conserved region is highlighted in yellow, positively charged region in blue and C-terminal PBS binding region in purple. The conserved FxxM motif is highlighted with a dashed box. **(D)** AlphaFold predicted structure (colored according to prediction confidence) of residues 1-121 of ApcG (not present in the cryoEM structure) of the combined with model of the PBS interaction domain from the cryo-EM structure (purple). Phosphorylation sites are shown as red spheres.

### The phosphorylation status of ApcG impacts growth and activity of photosystems in *Synechocystis*

In order to understand the physiological role of ApcG (*sll1873* gene locus), we generated a deletion strain replacing the native coding sequence of *apcG* by a chloramphenicol resistance cassette (Δ*apcG)*. Additionally, we complemented this deletion strain by replacing the native *psbA2* gene copy (Englund et al., 2016) with the *apcG* wild type open reading frame, and two phospho-mimicking versions to test the impact of the phosphorylation sites in the positively charged domain (**Supp. Figure S1**). This strategy ensures strong expression of the genes under the promoter of *psbA2* (P*psba2*) in *Synechocystis*. The generation of ApcG versions for testing the impact of its phosphorylation sites was achieved by mutating codons coding the residues 46-48 from *apcG* to glutamic acid (phospho-mimicking, TTS/EEE) or to alanine (permanent non-phosphorylated, TTS/AAA). Growth of wild type and *apcG* deletion strains showed no differences under normal conditions (constant light, 30 μmol photons m^−2^·s^−1^) as well as under stress conditions such as high light (400 μmol photons m^−2^·s^−1^ light intensity) and light to dark intervals (12 h light and 12 h darkness; **Supp. Figure S2**). However, when the deletion strain was grown under light-limiting conditions (10 μmol photons m^−2^·s^−1^), its growth was strongly impaired compared to the wild type (**Figure 2A**). Furthermore, when complemented with the phospho-mimicking *apcG^TTS/EEE^* growth was delayed during exponential phase under both 10 and 25 μmol photons m^−2^·s^−1^ compared to the deletion mutant and the other complementation strains (**Figure 2A**). Additionally, comparison of the strain’s growth using blue light (450 nm), green light (530 nm), red light (615 nm, exciting PBS) and far-red light (730 nm, exciting PSI) (Fuente et al., 2021) showed that the phospho-mimicking *apcG^TTS/EEE^* strain only shows reduction in growth when using red and far-red light, which could indicate an imbalance in the activity of the photosystems (**Supp. Figure S3**). Whole-cell absorption spectra of wild type and *apcG* deletion strains cultivated under normal conditions (i.e., white light of intensity 25-30 μmol photons m^−2^·s^−1^) showed no differences (**Figure 2B**), while the phospho-mimicking version had a distinct increase in absorption around 628 nm (**Figure 2C**). Analyses of the pigments present in the strains revealed that the phospho-mimicking strain contains a higher ratio of carotenoids to chlorophyll yet no difference in phycobiliprotein content compared to wild type (**Figure 2D**). Interestingly, the ratio of carotenoids and chlorophyll to culture turbidity, representing the amount of pigments per cell, was lower in the phospho-mimicking strain compared to wild type and the *apcG* deletion strain. The increase of carotenoids to chlorophyll ratio in the phospho-mimicking strain likely accounts for the spectral difference in whole-cell absorption (**Figure 2C**).

**Figure 2.**
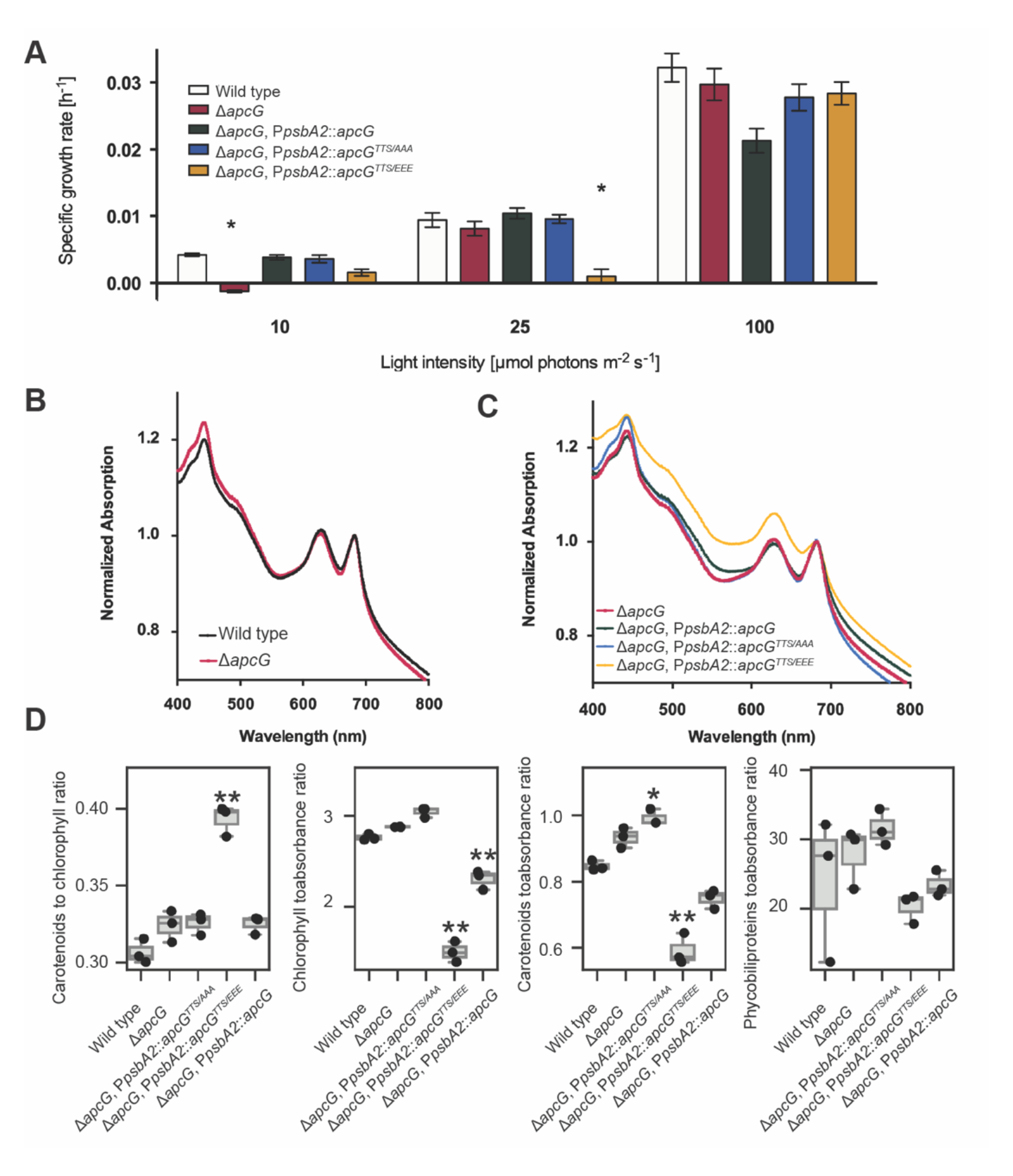
Impact on growth in *apcG* deletion strains. **(A)** Specific growth rate comparison of the *apcG* deletion, complementation strains and wild type under 10, 25 and 100 μmol photons m^−2^·s^−1^. Values represent means of four independent replicates ± SD and asterisks show statistical difference compared to wild type according to Student’s *t* test (two-sided, *P* < 0.05)**. (B)** Whole-cell absorption spectra for wild type *Synechocystis* and *apcG* deletion mutant, **(C)** and for *apcG* deletion and its complementation strains. Values correspond to averages of three biological replicates normalized to the second absorption peak of chlorophyll *a* at 680 nm. **(D)** Comparison of the relative pigment composition among *apcG* strains. Pigment concentrations were measured in three biological replicates in μg / mL units, and culture turbidity was measured as absorbance at 720 nm. A single asterisk represents statistical significance at *P* value of 0.05, and two asterisks represent statistical difference at *P* value of 0.01.

To gain further insight into the photosynthetic performance of the strains, low temperature (77 K) fluorescence spectra were recorded. In these experiments, we added an additional complementation strain using a construct coding for an *apcG* lacking the first 20 residues (of the N-terminal domain, **Figure 1B**). Excitation of chlorophyll at 430 nm revealed an increase of PSI fluorescence at 728 nm in the *apcG* mutant compared to the wild type. An even higher PSI fluorescence was observed in the complementation strains using the truncated *apcG^Δ1-20^* and the phospho-mimicking *apcG^TTS/EEE^* compared to wild type and non-phosphorylated strains (**Figure 3A**). Interestingly, when PBSs were excited at 590 nm, the phospho-mimicking strain continued to show a higher PSI fluorescence while exhibiting a decreased PSII fluorescence (**Figure 3B**). Other strains show similar behavior to the wild type when exciting PBS at 590 nm apart from *apcG* deletion strain that showed a mild increase in PSI fluorescence compared to wild type (**Figure 3B**).

**Figure 3.**
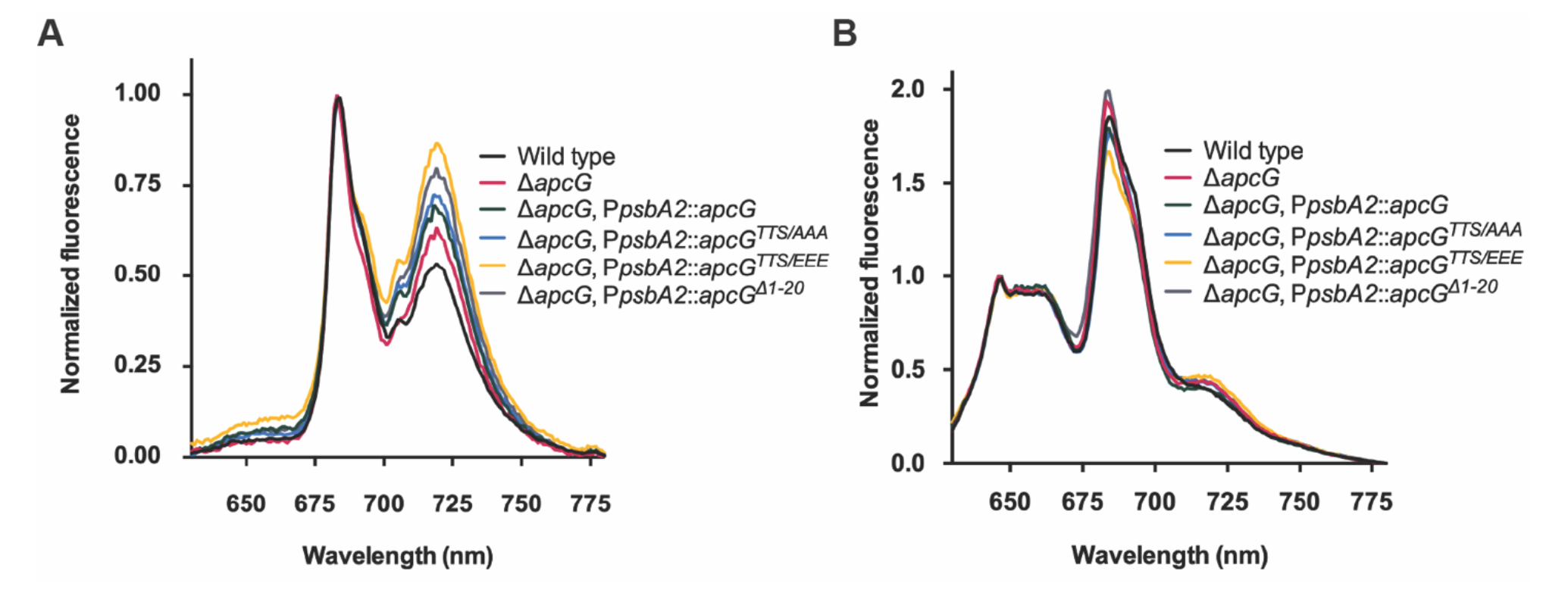
*apcG* deletion and phosphorylation impacts PSII and PSI energy balance. *Synechocystis* cultures were grown to compare their fluorescence under low temperature (77 K). Each curve on the spectra represents the mean from three biological replicates. All cultures were pre-incubated at room temperature in darkness for 20 min before measurements. **(A)** Fluorescence emission spectra of chlorophyll from *Synechocystis* with an excitation of 430 nm. The first peak at 680 nm corresponds to the fluorescence from PSII while the one around 720 nm to PSI. Values correspond to mean of three biological replicates while normalizing the data to the peak of PSII at 680 nm. **(B)** Fluorescence emission spectra of *Synechocystis* strains by exciting PBS at 590 nm. The fluorescence peak at 650-660 nm correspond to PBS, 680 nm to PSII and 720 nm to PSI. Values correspond to the means of three biological replicates while normalizing the data to the peak of PBS at 646 nm.

Since the *apcG* deletion, as well as the complementation strains with *apcG^Δ1-20^* and phospho-mimicking *apcG^TTS/EEE^* showed striking increase in PSI fluorescence at 77 K, we investigated the possible role of ApcG in state transitions. State I was induced in all tested *Synechosystis* strains by pre-treating the cultures with blue light, and state II was induced by pre-incubating the cultures in darkness. To monitor the differences in each state, 77 K fluorescence spectra of the cultures were recorded by exciting PBS at 590 nm. Surprisingly, all strains showed the same behavior as wild type under state I or II (**Supp. Figure S4**), indicating that the imbalance of PSI and PSII observed by exciting chlorophyll *a* (**Figure 3A**) is not related to state transitions.

Because the phospho-mimicking complementation strain showed higher PSI fluorescence, we analyzed the steady-state levels of proteins from PBS, PSII and PSI. Interestingly, immunoblots revealed no differences between strains in the levels of marker proteins for PSI (PsaB), PSII (PsbA) and PBS (APC) (**Figure 4A**). In order to discard the possibility that changes in the composition of thylakoid super-complexes could account for the higher PSI fluorescence observed in the *apcG* deletion strain, we compared the thylakoid super-complexes by blue native gels, which showed no differences between wild type and the *apcG* deletion strain (**Figure 4B**). Additionally, isolated PBS from wild type, *apcG* deletion and its complementation strains showed that the absence of ApcG did not impact the absorbance or fluorescence spectra of PBSs (**Supp. Figure S5**). Overall, our physiological comparison of *Synechocystis* strains shows that PBSs enhance energy transfer towards PSI when ApcG is truncated or phosphorylated, indicating a regulatory role impacting photosynthesis without disturbing the native organization of thylakoid super-complexes.

**Figure 4.**
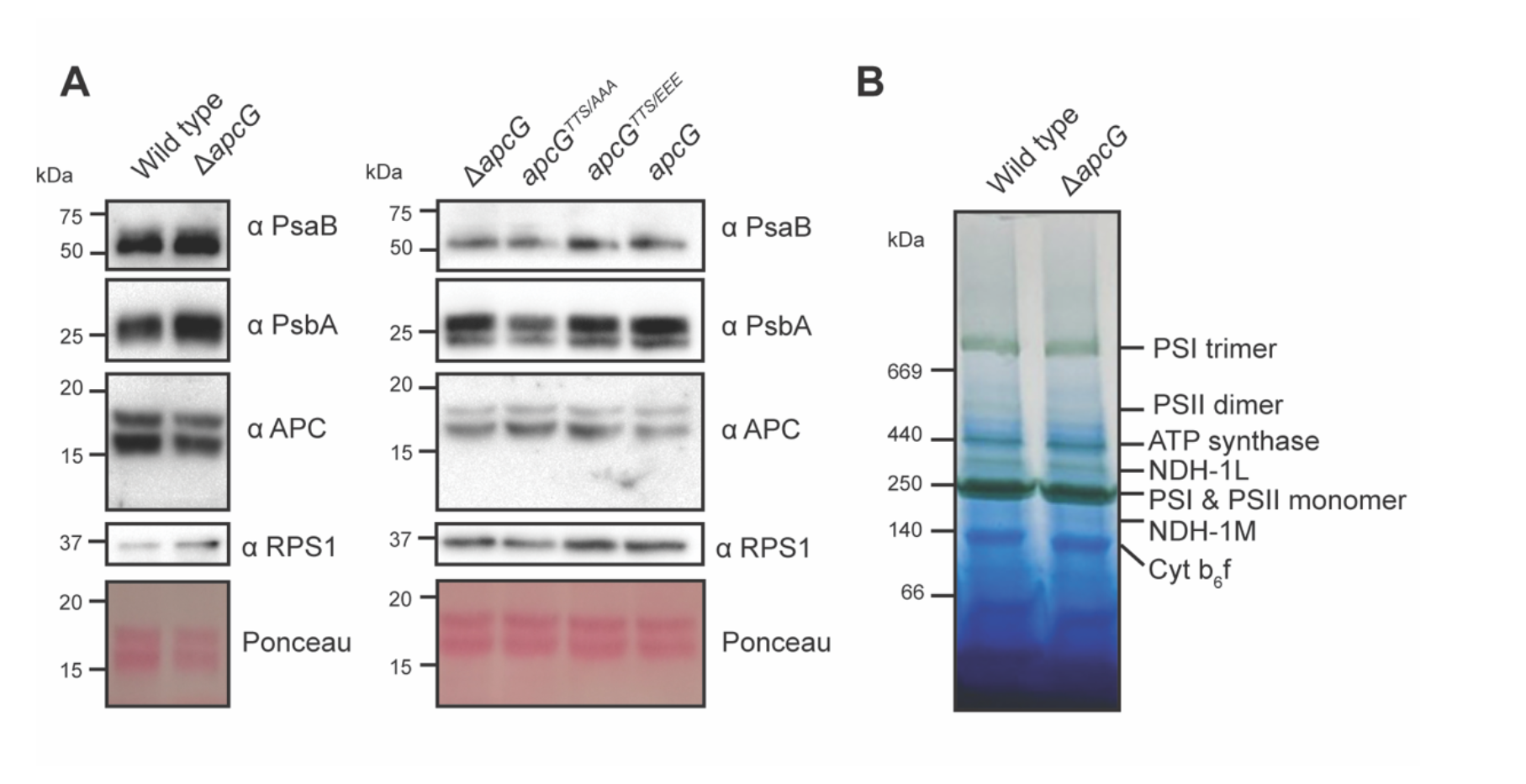
Impact of ApcG deletion on thylakoid super-complexes in *Synechocystis*. **(A)** Whole-cell protein extracts were obtained from *Synechocystis* strains to compare the accumulation of marker proteins for PBS (APC), PSI (PsaB) and PSII (PsbA). The ribosomal protein RPS1 was used as loading control. A total of 20 μg of protein were loaded for each lane. A representative immunoblot out of three biological replicates is shown. **(B)** Thylakoid membrane fractions from wild type and *apcG* deletion mutant were solubilized and separated on blue native gels to analyze the accumulation of the major thylakoid super-complexes. A total of 30 μg of chlorophyll were loaded for each strain. A representative native gel from one of three biological replicates is shown.

### The N-terminal domain of ApcG binds PSII

The structure of *Synechocystis* PBS shows that the C-terminal domain of ApcG binds to one of the two bottom core cylinders but the N-terminal and middle domains are not resolved (Dominguez-Martin et al., 2022). To investigate interaction partners of the N-terminal portion, we designed a version of ApcG where the PBS-binding domain is replaced by a His-tag (**Figure 5A**). This allowed both the rapid purification of the recombinant protein from *E. coli* as well as performing pull-down experiments using nickel affinity resin. We incubated *in vitro* ApcG bound to nickel resin with solubilized thylakoid membranes and noticed that the eluate after washing was green. This was not observed in the negative control (no ApcG). Immunoblot analyses to detect marker proteins for PSI and PSII showed that proteins from PSII were pulled down with the truncated ApcG (**Figure 5B**). The eluate was then separated on clear native (CN) gels showing the presence of one specific super-complex (**Figure 5C**). A second dimension of SDS-PAGE from the CN gel suggests that the super-complex found could correspond to PSII when compared to thylakoid super-complexes (**Figure 5D**). Indeed, immunoblots analyses of the CN gels using antibodies against PsbA showed that it corresponds to PSII (**Figure 5E**). Additionally, immunoblots using antibodies against the His-tag showed the presence of ApcG co-migrating with the PSII super-complex, confirming their interaction (**Figure 5E**).

**Figure 5.**
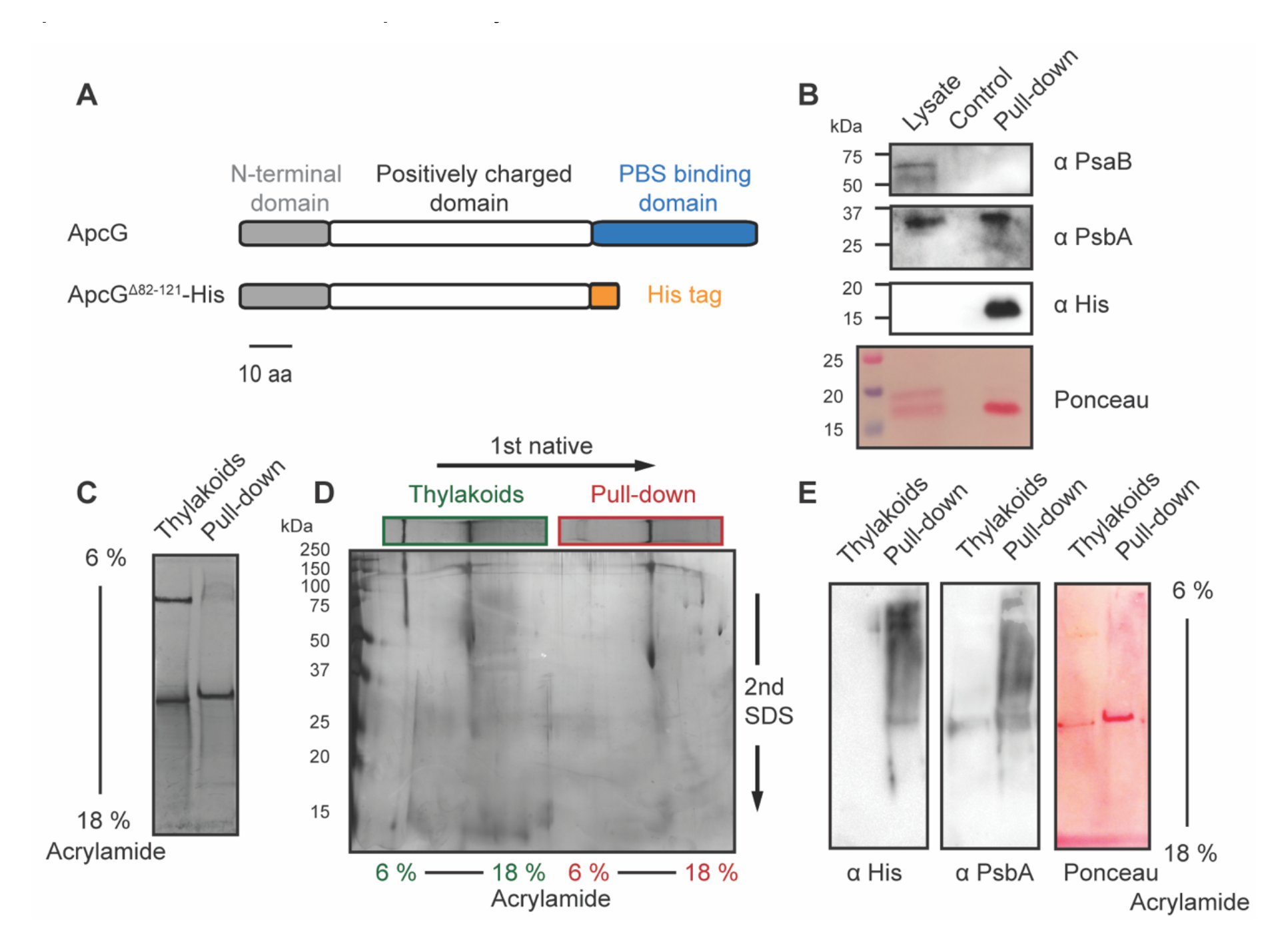
The N-terminal domain of ApcG interacts specifically with PSII. **(A)** Schematic representation of ApcG domains and the ApcG truncation used for pull-down experiments. **(B)** Immunoblots of the pull-down eluate using antibodies to detect the presence of PSI (anti-PsaB) and PSII (anti-PsbA). Whole-cell protein extract was used as a positive control for marker proteins. **(C)** Solubilized thylakoid membranes were incubated with nickel beads containing ApcG^Δ82-121^-His. After washing the beads and eluting with 200 mM imidazole, the eluate was separated in CN gels. For comparison, solubilized thylakoid membranes were loaded alongside the eluate from the pull-down experiment. A total of 1.3 μg of chlorophyll was loaded into each lane. **(D)** Second dimension of CN gel lanes by SDS-PAGE and stained with silver staining. **(E)** Immunoblots of the first CN gel dimension of pull-down eluate with total thylakoid super-complexes. Antibodies against His-tag were used to detect the presence of ApcG^Δ82-121^-His. Additionally, antibodies against PsbA were used as a marker for PSII. Images from section (**B**) to (**E**) correspond to a representative experiment from three biological replicates.

Our results using complementation strains with the phospho-mimicking version of *apcG* suggest that phosphorylation of the positively charged domain of ApcG could impact its interaction with the thylakoid super-complexes or the membrane. In order to assess this possibility, we generated two additional versions of the truncated ApcG by mutating the same residues used for the complementation strains, i.e., a permanent non-phosphorylated (TTS/AAA) and a phospho-mimicking version (TTS/EEE). Pull-down experiments showed a decrease of PSII proteins pulled down by the phospho-mimicking version while wild type and the non-phosphorylated version showed a similar amount of proteins from PSII (PsbA) being pulled down (**Figure 6A**). Furthermore, when using three times the amount of phospho-mimicking (TTS/EEE) truncated ApcG for the pull-down experiment relative to wild type and the permanent non-phosphorylated (TTS/AAA), similar amounts of PsbA were pulled down. This implies that the ability of ApcG to interact with PSII in the phospho-mimicking version (TTS/EEE) is reduced compared to the wild type ApcG. The PSI core protein PsaB was not detected in any of the pull-downs, indicating that neither wild type ApcG nor any of the phospho-mimicking mutants interact with PSI (**Figure 6B**). In order to corroborate that the proteins pulled down corresponded to the super-complex of PSII, CN gels were run using the same amount of protein loaded in each lane and blotted to a membrane to detect PSII super-complexes using antibodies against PsbA. Indeed, all three ApcG versions were able to pull down PSII super-complexes (**Figure 6C**). These results indicate that the N-terminal region of ApcG specifically interacts with PSII while the phosphorylation of ApcG in its positively charge middle domain impairs the ApcG-PSII interaction. Additionally, the phosphorylation of ApcG does not alter its specificity for PSII.

**Figure 6.**
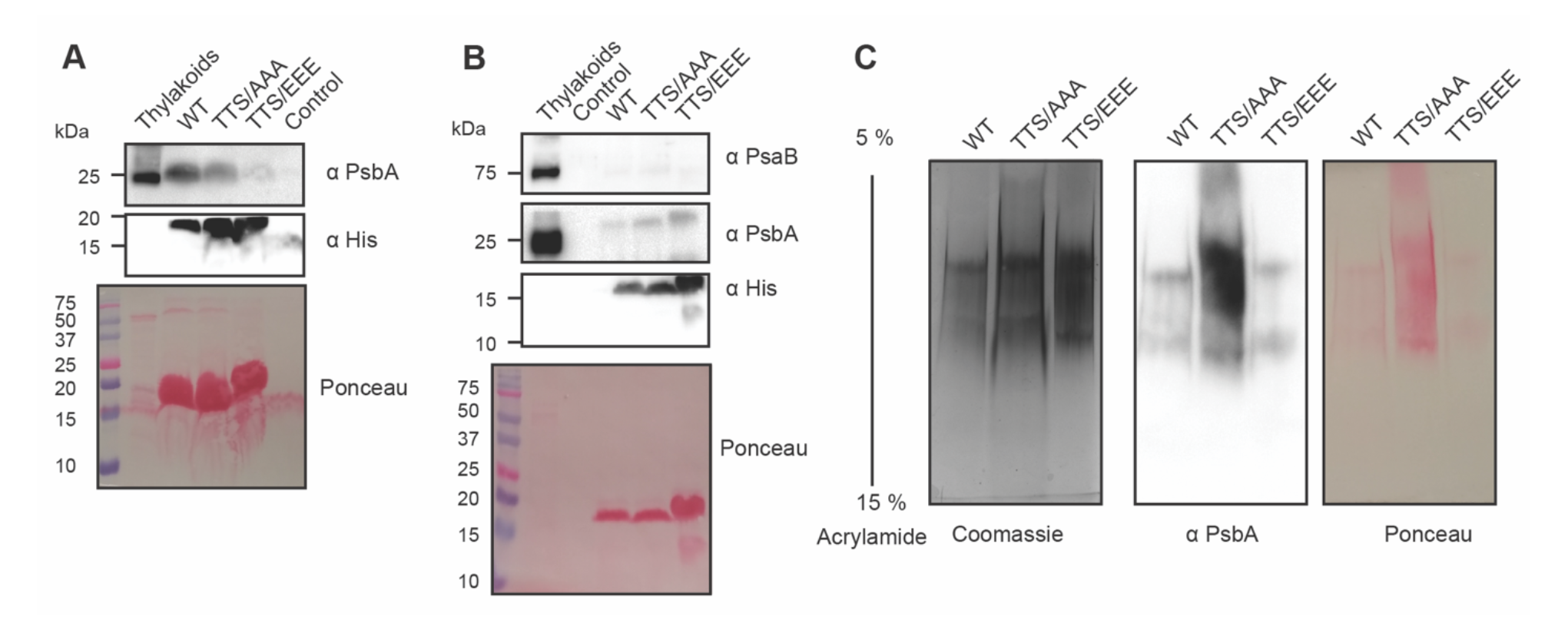
The phosphorylation status of ApcG influences its interaction with PSII. Pull-down experiments using the truncated form of ApcG with a mutated version for a permanent non-phosphorylated (TTS/AAA) and a phospho-mimicking version (TTS/EEE). Solubilized thylakoid membranes from the ApcG deletion strain were used to perform pull-down experiments. A representative experiment (one of four) is shown. (**A**) Western blot analysis for the detection of PSII marker protein PsbA pulled down by each ApcG version used. Anti-His antibodies were used to detect the three ApcG His-tagged versions. The experiment was performed using the same amount of truncated ApcG versions and solubilized thylakoids. (**B**) Pull-down experiment using three times more of the phospho-mimicking version (TTS/EEE) compared to wild type and permanent non-phosphorylated (TTS/AAA). Antibodies against marker proteins for PSII (PsbA) and PSI (PsaB) were used to detect the super-complexes pulled down by each truncated form of ApcG. **(C)** Native gel analysis for the pull down of PSII super-complexes (2 μg of protein loaded per lane). A Coomassie stained gel is shown as well as the immunoblot detection of PSII marker protein PsbA. Images from section (**A**) to (**C**) show a representative experiment from three biological replicates.

## DISCUSSION

In cyanobacteria, the energy transfer from PBS to the photosystems is known to have several modes of regulation. For example, in response to high light there is the PBS quenching mechanism mediated by the orange carotenoid protein, which dissipates the excess of energy absorbed by PBS (Dominguez-Martin et al., 2022, Leverenz et al., 2015, Kerfeld et al., 2017) thus reducing the likelihood of saturating the electron transport chain in the thylakoid membrane. Likewise, as a response to differences in light quality, the PBS can undergo state transitions by shifting between state I (PSII saturation or reduced plastoquinone pool) and state II (PSI saturation or oxidized plastoquinone pool) to balance the activity of both photosystems and prevent the accumulation of reduced intermediates in the electron transport chain (Emlyn-Jones et al., 1999, Xu et al., 2012, Bonaventura and Myers, 1969, Hodges and Barber, 1983). Cyanobacterial state transitions, unlike those of plants, is not well understood. Several models have been put forward to explain the mechanism behind this regulatory process. For example, the “attachment/detachment” model in which PBS are released from PSII (state I) and interact with PSI (state II) (Sarcina et al., 2001, Mullineaux et al., 1997, Yang et al., 2007). Another model is the “spillover” mechanism in which PBS maintain their interaction with PSII, but excitation energy spills over from PSII to PSI (McConnell et al., 2002, Folea et al., 2008, Olive et al., 1997).

Several of our observations are consistent with a spill over mechanism. Our studies using deletion and complementation strains for different version of ApcG showed that this linker protein is indeed necessary for the efficient transfer of energy to drive photosynthesis, especially under light-limiting conditions (10 μmol photons m^−2^ s^−1^ light intensity). Furthermore, under normal conditions (25 μmol photons m^−2^ s^−1^ light intensity), the phospho-mimicking *apcG^TTS/EEE^* strain showed delayed growth compared to all others including the deletion *apcG* strain (**Figure 2A**). Analyses of pigment content showed an increase in carotenoid per chlorophyll ratio in the phospho-mimicking *apcG^TTS/EEE^* strain (**Figure 2C**). Similarly, deletion mutants for proteins affecting photosynthetic function in *Synechocystis* showed that indeed a higher carotenoid per chlorophyll ratio is symptomatic of light stress with increased anti-oxidant compounds content such as myxoxanthophyll and zeaxanthin (Cunningham et al., 2010, Havaux et al., 2003, Maeda et al., 2005, Schafer et al., 2005). Strikingly, low temperature fluorescence spectra of these strains displayed an increase in PSI fluorescence when exciting chlorophyll with the highest one observed in the *apcG^Δ1-20^*and phospho-mimicking *apcG^TTS/EEE^* strains. This phenomenon was observed in both the raw and normalized spectra among strains. (**Figure 3A**). Considering that (i) in these strains PBSs impact the PSI fluorescence even when PBSs are not absorbing energy (**Figure 3A**) and (ii) among strains, there is no difference in PSI accumulation or super-complex distribution to account for the differences in PSI fluorescence (**Figure 4A**), the mechanism that best accounts for these observations is “spillover”. Furthermore, when observing low temperature fluorescence spectra exciting PBS at 590 nm, a similar pattern is observed with a higher PSI fluorescence (720 nm) in the phospho-mimicking *apcG^TTS/EEE^*strain and a decrease in PSII fluorescence (680 nm) (**Figure 3B**). Thus, as a consequence of the truncation or phosphorylation of ApcG, our results support an indirect effect of the PBS linker protein ApcG on the transfer of energy from PSII towards PSI most likely via “spillover”. The spillover model has generally been associated with state transitions (McConnell et al., 2002, Li et al., 2006, Li et al., 2004). However, our results show that the putative spillover involving ApcG does not play a role in state transitions because all the mutant strains used in this study are able to transition between state I and II, comparable to wild type (**Supp. Figure S4**). Therefore, the spillover phenomenon observed in the *apcG* deletion as well as in the phospho-mimicking *apcG^TTS/EEE^* strains implies that this process is not related to state transitions. Nonetheless, these two mechanisms (spillover and attachment/detachment) could very well happen in parallel, as it has been suggested (Mullineaux et al., 1997, Mullineaux, 2014, McConnell et al., 2002, Li et al., 2004, Li et al., 2006). Thus, the phenotype observed under light-limiting conditions in the deletion *apcG* strain highlights the imbalance in PSI and PSII, leading to decrease in the linear electron transport.

The PBS linker protein ApcG offers an opportunity to investigate the interface between PBS and the membrane-embedded photosystems because of its position at the base of the two core cylinders (Dominguez-Martin et al., 2022). Due to its occurrence along with ApcE, which by its domain architecture defines the cylindrical core structure of PBS, the function of ApcG in PBS might be quite conserved among cyanobacteria. Hence, ApcG is expected to facilitate the interaction of PBS with super-complexes in the thylakoid membrane. Our pull-down experiments using a truncated form of ApcG lacking the PBS binding domain shows that the N-terminal region of the ApcG interacts with PSII from solubilized thylakoids, presumably via the conserved domains of the N-terminus or the positively charged middle region. Furthermore, in the ApcG^TTS/EEE^ phospho-mimicking mutant protein, the interaction with PSII is impaired. Additionally, the phospho-mimicking *apcG^TTS/EEE^*strain showed delayed growth (**Figure 2A**) suggesting that phosphorylation could as well cause repulsion of the middle domain from the negatively charged head groups of thylakoid lipids. Our results support a regulatory role for phosphorylation of ApcG by altering the charge of the positively charged middle domain, which influences its interaction with PSII (**Figure 6A**). Indeed, phosphorylation of other PBS proteins has been reported (Toyoshima et al., 2020), (including ApcA, ApcC, ApcF, CpcA, CpcB) with CpcB phosphorylation having a direct impact on state transitions (Chen et al., 2015). Interestingly, 77K spectra after excitation of the PBS at 590 nm showed no major differences in PSII maxima among strains (**Figure 3B**), implying that ApcG is not the only protein involved in the interaction between PBS and PSII. This is supported by a red alga PBS-PSII structure that found three unknown connector proteins in PBS (Li et al., 2021). As observed in low temperature fluorescence spectra, PSI receives more energy in the *apcG* deletion, *apcG^Δ1-20^*and phospho-mimicking *apcG^TTS/EEE^* strains, yet pull-down experiments showed no accumulation of PsaB (a core protein of PSI) (**Figure 6B**). These results suggest that, even though in these strains PSI receives more energy compared to wild type, this phenomenon occurs without a direct physical interaction between ApcG and PSI. Therefore, the effect that ApcG exerts on the fluorescence of PSI might be the result of PSII-PSI interaction allowing the energy from PSII to spill over to PSI. Indeed, there is experimental evidence for super-complexes involving PBS-PSII-PSI as well as PSII-PSI except for PBS-PSI (Liu et al., 2013, Beckova et al., 2017, You et al., 2023). A super-complex of PBS-PSII-PSI would allow spillover to occur; we observe evidence for ApcG interacting solely with PSII yet affecting the energy transfer from PSII to PSI. A recent PBS-PSII-PSI super-complex structure has been reported in red algae *Porphyridium purpureum*; an ApcG homolog (L_pp_2) is found in the super-complex interacting with a PSII dimer. In contrast, the PSII-PSI interaction does not involve L_pp_2 (You et al., 2023), consistent with our observations (**Figure 3A and 6B**).

Our experiments explored the influence of phosphorylation using only ApcG point mutations. Under similar growth conditions other proteins of a PBS-PSII complex might also be phosphorylated. Indeed, PSII undergoes phosphorylation in several subunits (PsbA, PsbB and PsbC) as well as for PSI (PsaA, PsaB, PsaC, PsaD, PsaE, PsaF, PsaL) (Toyoshima et al., 2020, Angeleri et al., 2016). Thus, our results cannot exclude that ApcG interacts with PSI or a PSI-PSII super-complex under conditions in which ApcG, as well as subunits of PSI and PSII, undergo phosphorylation.

Collectively, our results show that the linker protein ApcG is necessary for efficient light harvesting under light limiting conditions. While the C-terminal PBS-binding domain binds to each of the two core bottom cylinders (Dominguez-Martin et al., 2022), its N-terminal region interacts specifically with PSII as shown in our *in-vitro* pull down assays. Furthermore, when phosphorylated in its positively charged middle domain, the interaction of ApcG with PSII is hindered, which is correlated with a slower growth rate (**Figure 2A and 6A**). The truncation as well as the phosphorylation of ApcG in complementation strains showed an increase of energy absorbed by PSI even under conditions where PBS do not absorb light, suggesting a role for ApcG in the spillover from PSII to PSI.

## MATERIALS AND METHODS

### Cyanobacteria growth conditions

*Synechocystis* sp. PCC 6803 strains were grown in BG-11 medium (Rippka et al., 1979), buffered to pH 8 with 10 mM HEPES, at 28–30°C under constant light (25– 30 μmol photons m^−2^ s^−1^) and enriched with 3% CO_2_, unless otherwise stated, in shaken liquid cultures (160 rpm). For selection of mutants on plates, BG-11 containing 3 g L^−1^ sodium thiosulfate was solidified with 1.2% Difco agar. Antibiotic concentrations used for selection of *Synechocystis* mutants were chloramphenicol at 25 μg ml^−1^, or spectinomycin at 20 μg ml^−1^.

Cyanobacteria growth under these conditions (hereafter referred to as normal conditions) was compared by cultivating the strains in flasks in batch regime starting with OD_750_ 0.05. To apply light stress, the multi-cultivators (MC 1000-OD, Photon System Instruments, PSI, Czech Republic) were used for cultivations under constant high light (400 μmol photons m^−2^ s^−1^) or for 12 hours darkness followed by 12 hours illumination (30 μmol photons m^−2^ s^−1^) for a period of 10 days.

The strains were further cultivated in Multi-cultivators MC-1000-MIX (Photon System Instruments) in turbidostat regime, in BG-11 cultivation medium as described in the previous section at 30 °C and at 0.5% CO_2_. The strains were cultivated under 10, 25 and 100 μmol photons m^-2^ s^-1^ of warm white as well as green, red and far-red light under 25 and 100 μmol photons m^-2^ s^-1^. The illumination was provided by LEDs with the following peaks and half-bandwidths: blue: 450 ± 25 nm, green: 537 ± 40 nm, red: 615 ± 25 nm and far-red: 730 ± 15 nm (**Supp. Figure S6**). The density range of the turbidostat cultivation was set to OD_720_ 0.5 – 0.51, (approximately 10^7^ cells mL^-1^). The cultures were cultivated under each light condition for at least 22 h, to secure full metabolic acclimation (Zavrel et al., 2019). After the acclimation period, specific growth rates were estimated from the change of OD_720_.

### Generation of *Synechocystis* deletion and complementation strains

The generation of deletion strain for *apcG* was done by amplifying the 600 base pairs upstream as well as downstream its gene locus (Sll1873) using primers oREC1 (5’- CCCTCAAACCCCAAACGATT) and oREC2 (5’-CGGGGCGAATGGTTTCTAAC) and cloning into pJET1.2. The *apcG* gene was then replaced by a SacI site through inverse-PCR using primers oREC3 (5’-GAGCTCTTTAATGTGGTTCTCCTAATTG) and oREC4 (5’-AAACCTCATTGATTTACTGTTTTATAC). This construct was used to introduce an insert from pRL1075 compatible for bacterial conjugation and containing chloramphenicol resistance cassette (Black et al., 1993). The insert was introduced by digestion and ligation using SacI, resulting in the construct pSL399 for transformation of wild type *Synechocystis*. For complementation strains, the open reading frame from *apcG* was amplified with oREC9 (5’-TCGTCATATGTTAAAAAAATTGTTTGGCGCT) and oREC10 (5’-GTGCTCGAGACCGGAGCGTTTAACCTTAACTTGGCGAG) and cloned into pET-28a(+) using restriction digestion enzymes BamHI and XhoI, resulting into the construct pREC4. A region containing *apcG* from pREC4 was amplified using oREC11 (5’- CCAATCCGGAGGATCCTATAGTTCCTCCTTTCAGCAA) and CK10 (5’-TAATACGACTCACTATAGGG) and cloned into pPSBA2KS (Lagarde et al., 2000) using restriction digestion enzymes BamHI and NdeI, resulting in the plasmid pREC6. A Bom site compatible for bacterial conjugation was incorporated into pREC6 by inverse-PCR and ligation using oREC25 (5’- CACTCTCAGTACAATCTGCTCTGATGCCGCATCGAGCTCTGTACATGTCCGCGG) and oREC26 (5’-CACCATATGCGGTGTGAAATACCGCACAGATGAGAAGTACTAGTGGCCACGTGG) resulting in plasmid pREC9. A spectinomycin resistance cassette was incorporated into pREC9 by amplifying the *aadA* gene from pRL3332 (Nieves-Morion et al., 2017) using oREC46 and oREC47 having BamHI at both extremes that was used to clone it into pREC9 using its unique BamHI site, resulting in plasmid pREC17. Finally, the C-terminal His tag from pREC17 was removed by inverse-PCR using oREC50 (5’- TGAGATCCGGCTGCTAACAAAG) and oREC51 (5’-GCGTTTAACCTTAACTTGGCGAGCCA) resulting in pREC28 bearing a spectinomycin cassette for selection in *Synechocystis* and a Bom site for bacterial conjugation of the deletion strain for *apcG*. The phospho-mimicking complementation constructs were generated by inverse-PCR using pREC28 as template with primers oREC38 (5’- CCGGCTCCGGCTGCTGCTAAAAAAACT) and oREC39 (5’-TTTTTCCACCGGAGCTACCTCCG) for the permanent non-phosphorylated version (residues 46-48 TTS into AAA, construct pREC30) and the phospho-mimicking version (residues 46-48 TTS to EEE, construct pREC31) using primers oREC39 and oREC42 (5’- CCGGCTCCGGAAGAAGAAAAAAAAACT).

Furthermore, constructs for over-expression in *E. coli* for a truncated version of ApcG removing its PBS binding domain (residues 82 to 121) and keeping a C-terminal His tag was obtained by inverse-PCR using pREC4 as template and primers oREC32 (5’- AACTTTGGCCTTGGGAGCCGGGG) and oREC33 (5’-GGTCTCGAGCACCACCACCAC), to then sub-clone this region using BamHI and NdeI into pBF6 resulting in plasmid pREC54 for tetracycline bacterial induction. Likewise, phosho-mimicking versions for over-expression in *E. coli* were obtained by inverse-PCR using pREC54 as template and primers oREC38 and oREC39 (permanent non-phosphorylated, pREC56) as well as oREC39 and oREC42 (phospho-mimicking version, pREC55). Transformation of wild type as well as the ApcG deletion mutant was performed by bacterial conjugation as described by Black et al. (1993).

### *Synechocystis* strains genotyping

Cyanobacteria strain cultures were grown to OD_750_ 0.6 – 0.8 under normal conditions and their genomic DNA was extracted by the phenol - chloroform method described by Billi et al. (1998). For amplification of the wild type allele the PCR used primers oREC12 (5’- AGACGGGGAAAAGGCTCTAC) and oREC13 (5’- CCGCTTCAATTTCCTCGTCC). However, the deletion allele was amplified with primers oREC12 and oREC27 (5’- TTCCACGGACTATAGACTATACT). The over-expression insertion was detected by amplifying a fragment using primers oREC57 (5’- CCCAGGGACAATGTGACCAAAAAATTCA) and oCK11 (5’- GCTAGTTATTGCTCAGCGG).

### Protein expression and purification

Plasmids carrying the truncated form of ApcG for pull-down experiments (pREC54, pREC55 and pREC56) were transformed into *E. coli* BL21 DE3 (Invitrogen, Carlsbad, CA, USA). Cell cultures were grown in luria broth under 37°C till they reached OD_600_ ∼ 0.7, followed by induction with 10 μg ml^-1^ anhydrous tetracycline at 25°C overnight. One-liter cultures were centrifuged and resuspended in Buffer A (50 mM Tris pH 8, 200 mM NaCl) with protease inhibitor cocktail (Sigma, St. Louis, MO, USA), DNase I (Sigma) and lysed using two passes through a cell disruptor (Constant Systems, Aberdeenshire, UK) at 15 kPSI. The soluble fraction of the lysed sample was obtained by centrifugation for 30 min at 30,000 x g and 4°C. The recombinant proteins were purified loading the cell lysate supernatant to a 5 ml HisTrap HP column (GE Healthcare, Little Chalfont, UK), washed with Buffer A, followed by a 5-column volume (CV) of 90% Buffer A and 10% Buffer B (50 mM Tris pH 8, 200 mM NaCl, 500 mM imidazole) and eluted with a 5 CV gradient from 10 to 100% Buffer B. The recombinant proteins were further purified by cation exchange. The eluate from HisTrap was diluted with 50 mM Tris pH 8 10 times to reach 20 mM NaCl and loaded into pre-equilibrated cation exchange resin (TOYOPEARL SP- 650, column volume 5 ml) and performed the chromatography by gravity at 4°C. The column was then washed with 10 CV of buffer W (50 mM Tris pH 8, 20 mM NaCl), 5 CV with W2 (50 mM Tris pH 8, 50 mM NaCl) and eluted with buffer A. When purifying the truncated form of ApcG with residues 46-48 TSS mutated to EEE (phospho-mimicking), cation exchange step was omitted due to their weak binding to the resin. Protein concentration was measured using BCA method (Pierce BCA Protein Assay Kit, 23227, Thermo Scientific).

### Pull-down experiments with solubilized thylakoid membranes from *Synechocystis*

Cyanobacterial cultures of the *apcG* deletion strain grown for one week under normal conditions were collected by centrifugation and resuspended in 0.1 M phosphate buffer and pH 7.5. The cells were broken by French pressing, and the membranes were separated from the soluble proteins by centrifugating the sample for 30 min and 45,000 x g at 4°C. The thylakoids in the pellet fraction were resuspended in 10 ml of solubilization buffer (1 % dodecyl-beta-D-maltoside, 750 mM aminocaproic acid, 50 mM Bis-Tris pH 7 and 50 mM imidazole) and were incubated on ice for 30 min. After this the sample was centrifuged for 30 min at 30,000 x g at 4°C to discard insoluble membrane complexes. The soluble fraction corresponds to the solubilized thylakoid super-complexes whose protein and chlorophyll contents were quantified by BCA method and methanol extraction respectively. The solubilized thylakoid super-complexes were loaded into NTA nickel beads (0.8 ml column volume) pre-incubated with the truncated ApcG with His tag at its C-terminus in solubilization buffer with 50 mM imidazole. Beads were incubated under rotation for 1 hour at 4°C. The beads were then centrifuged for 2 min at 100 x g and the supernatant discarded to wash the beads 4 times with 10 CV of solubilization buffer and 50 mM imidazole. The elution was performed with 1.5 ml of solubilization buffer and 200 mM imidazole.

### Separation of super-complexes in first dimension native and second dimension denaturing gels

Solubilized thylakoid super-complexes as well as eluates from pull-down experiments were separated in native gels following the method described by Schagger and Vonjagow (1991). For clear native gels though, the same method described by Schagger and Vonjagow (1991) was followed but preparing the cathode running buffer as well as the sample loading buffer without Coomassie brilliant blue. After the separation of super-complexes in native gels, we further separated their protein content in a second dimension under denaturing conditions with 12% SDS-polyacrylamide gels supplemented with 4 M urea. The gels were stained using the method described by Blum et al. (1987).

### Preparation of total protein extracts

*Synechocystis* strains were grown in 10 ml BG-11 media in flasks of 25 ml under agitation under constant light (c. 25–30 μmol photons m^−2^ s^−1^) supplemented with 3% CO_2_ until they reached OD_750_ ∼ 1. Cultures were centrifuged and the supernatant discarded to resuspend cells in extraction buffer (50 mM HEPES pH 7.0, 25 mM CaCl_2_, 5 mM MgCl_2_, 10% [v/v] glycerol and protease inhibitor cocktail). Resuspended cells were broken by French pressing, and Triton X-100 was added to a final concentration of 1 % (v/v). After incubation on ice for 10 minutes, cell debris was discarded by centrifugation for 2 minutes at 2.000 g, 4°C and the supernatant rescued as total protein extract. Protein concentration was measured by BCA method.

### Cyanobacterial pigment analyses

To quantify chlorophyll and carotenoids, 1 mL culture was harvested in an Eppendorf tube. The cell pellet was suspended in 100% methanol, and absorption spectra of the extracted pigments were measured. The pigment concentration was calculated using calculations described by Zavřel (2015). Phycobiliproteins were quantified as described by Zavrel et al. (2018).

### Immunoblot analyses

Proteins separated into SDS-PAGE gels were transferred to a nitrocellulose membrane (Amersham^™^, Protran^®^). The membrane was blocked with 5% milk in TBS (Tris 20 mM and 150 mM NaCl) at room temperature for one hour then incubated with monospecific polyclonal antisera in TBS-T (Tris 20 mM, 150 mM NaCl and 0.01% tween-20) overnight at 4°C (anti-PsbA; AS05 084A; anti-PsaB; AS10 695; anti-APC; AS08 277; Agrisera, anti-His tag; TA150087; OriGene). The membrane was washed 3 times in TBST-T at room temperature for 15 minutes each wash followed by incubation with secondary polyclonal anti-rabbit antisera HRP for one hour at room temperature in TBS-T (Jackson ImmunoResearch, 111-035-003). After 3 additional washes with TBS-T, the membrane was visualized by the enhanced chemiluminescence technique.

### Isolation of PBSs from *Synechocystis*

Cyanobacteria cultures of one liter were grown under normal conditions for one week and harvested for resuspension in phosphate buffer (0.8 M, pH 7.5) supplemented with protease inhibitor cocktail (Sigma, St. Louis, MO, USA). Cells were broken by French pressing followed by the addition of 1% Triton X-100 and an incubation of 15 min at room temperature under darkness and gentle rotation. The soluble fraction containing PBSs was separated from the membrane fraction by centrifugation for 30 min at 30,000 x g and room temperature. The supernatant was rescued and centrifuged again for one hour at 42,000 x g and room temperature and the dark blue supernatant was separated from the green top with a syringe. These samples were loaded onto sucrose gradients composed of 1.5 M, 1 M, 0.75 M, 0.5 M and 0.25 M phases in phosphate buffer (0.8 M, pH 7.5), and separated by centrifugation at 25,000 rpm and room temperature overnight. Intact PBS fractions were recovered from the 0.75 M - 1 M interface of the sucrose gradients. The PBS protein content was measured by BCA method and proteins precipitated by trichloroacetic acid before being separated into SDS-PAGE.

### Measurements of absorption or fluorescence spectra

Whole-cells or PBS absorption spectra were recorded with a Varian Cary Bio 100 spectrophotometer (Agilent). Fluorescence spectra of isolated PBS samples were recorded with a fluorimeter (SpectraMax M2, Molecular Devices) exciting at 590 nm and emission spectra collected at room temperature from 610 to 800 nm.

### Fluorescence emission spectroscopy at 77 K

The cells were grown in BG-11 supplemented with 20 mM NaHCO_3_ and 10 mM HEPES-NaOH (pH 8.0) on a shaker at 30°C under constant light intensity at ∼50 µmol photons m^-2^ s^-1^. The cell concentration was adjusted to OD_630_ of 0.2-0.3 as measured by absorption spectroscopy using an integrating sphere (Shimadzu UV-3600i Plus with ISR- 603). The dark treatment was applied for 20 min to induce State II and subsequently, the blue light illumination was applied for 10 min to induce State I (Bhatti et al., 2020, Calzadilla and Kirilovsky, 2020, McConnell et al., 2002). Fluorescence emission was recorded at 77 K using a FluoroMax-4 spectrofluorometer (Horiba Scientific). The excitation wavelength was 430 nm or 590 nm with a 2-nm slit size. The emission wavelength measured was from 630 to 780 nm with a 2-nm slit size. Fluorescence emission for each sample was recorded consecutively three times to obtain averaged spectra. The results shown are averages of three independent biological replicates.

### Software

Figures were generated using Adobe Illustrator CS6. Graphs and statistical analyses were done using Python (Sanner, 1999) and GraphPad Prism version 6.0 (GraphPad Software, La Jolla, CA, USA) (www.graphpad.com). Structural figures were prepared with PyMOL (www.pymol.org). The sequence conservation logo was generated with Weblogo (Crooks et al., 2004).

## Abbreviations

PS: photosystem
PBS: phycobilisomes
APC: allophycocyanin

## Acknowledgments and funding

The research in the Kerfeld lab was supported by the Office of Science of the U.S. Department of Energy under award number DE-SC0020606. M.I. and K.K.N. were supported by the U.S. Department of Energy, Office of Science, Basic Energy Sciences, Chemical Sciences, Geosciences, and Biosciences Division under field work proposal 449B. K.K.N. is an investigator of the Howard Hughes Medical Institute.

## Author Contributions

R.E-C. designed and conducted the research, analyzed the data, and wrote the article, M.I. conducted low temperature fluorescence and analyzed the data, T.Z. conducted cyanobacteria growth experiments and analyzed data, S.L-Y. conducted pigment analyses of cyanobacteria strains and analyzed data, C.A.K. and M.S. designed research, analyzed the data and wrote the article. J.C., K.K.N. as well as all other authors provided comments on the manuscript and contributed to experimental design.

