## Supplemental data for "The phycobilisome linker protein ApcG interacts with photosystem II and regulates energy transfer to photosystem I in *Synechocystis sp.* PCC 6803"

**Abbreviations**: PS: photosystem; PBS: phycobilisomes; APC: allophycocyanin

### Supplemental data (6)

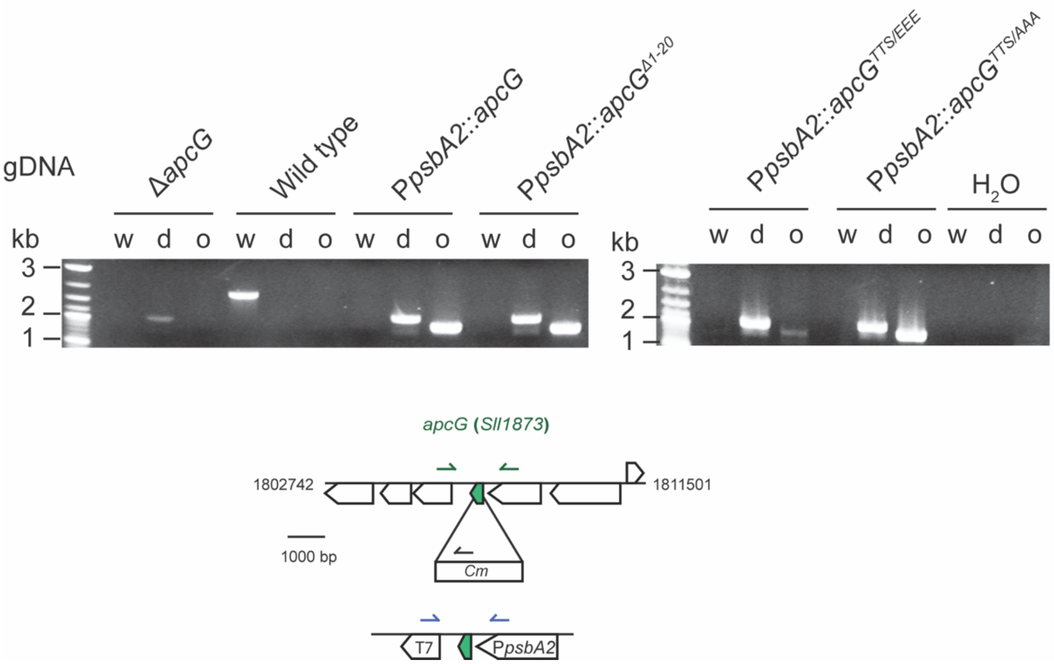


**Supplemental figure S1. Genotyping of deletion and complementation strains of *apcG*.** Genomic DNA (gDNA) was extracted from wild type, deletion, and the complementation strains to perform PCRs for the detection of the wild type allele (w), deletion cassette (d) or over-expression gene (o). In the lower panel it is shown the locus of *apcG* (*Sll1873*) as well as the primers used for each PCR reaction.

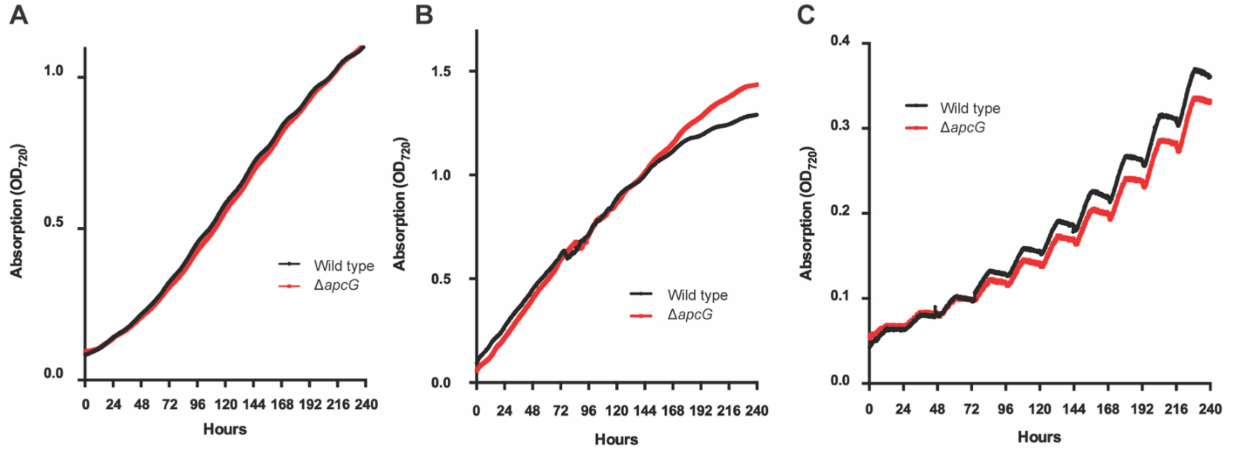


**Supplemental figure S2. Growth curve comparison of the wild type and *apcG* deletion strains under normal and light stress conditions.** Wild type and *apcG* deletion strains growth were compared under constant low light of 30 µmol photons $s^{-1} \cdot m^{-2}$ light intensity (**A**), constant high light of 400 µmol photons $s^{-1} \cdot m^{-2}$ light intensity **(B)** and light to dark periods of 12 hours each with 30 µmol photons $s^{-1} \cdot m^{-2}$ light intensity for the light periods **(C)**. Values correspond technical duplicates of a representative experiment from three biologically independent replicates. Growth curves were obtained with the instrument Multi-cultivator from PSI (MC 1000-OD) starting the cultures with 0.05 OD at 750 nm.

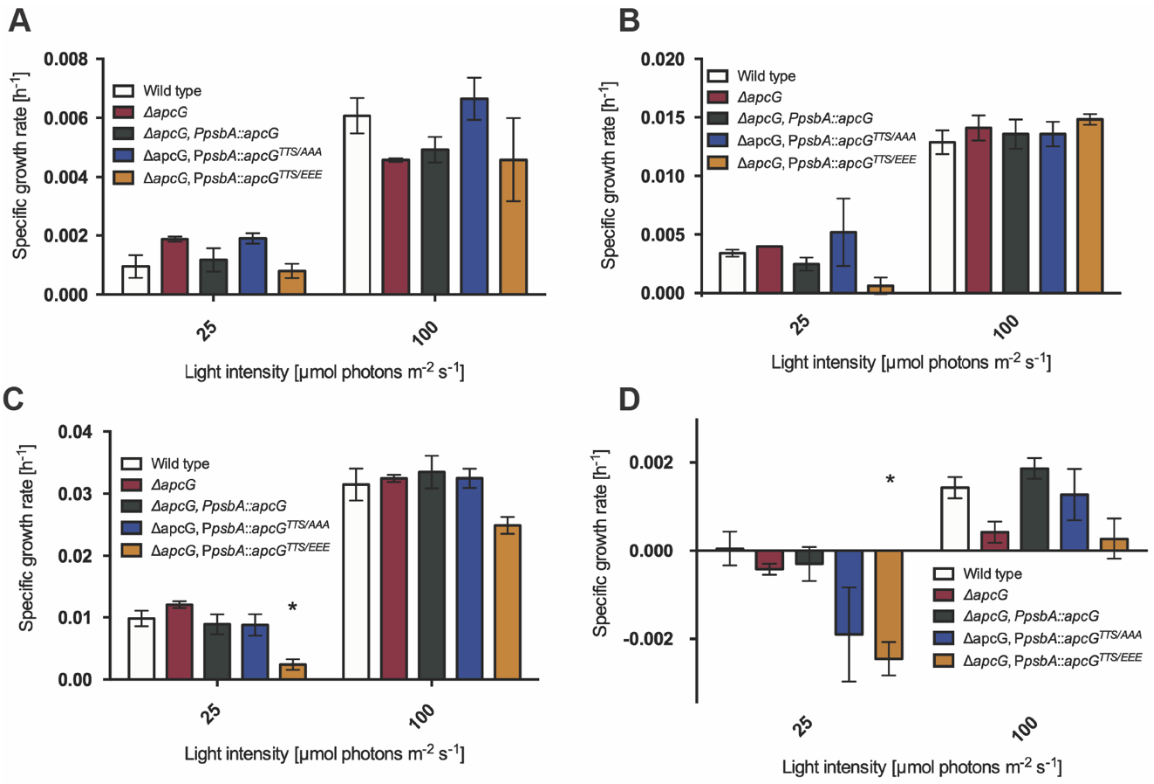


**Supplemental figure S3. Comparison of growth in *apcG* strains under different light qualities and quantities.** Cyanobacteria strains growth from *apcG* deletion, complementation strains and wild type was compared using blue light (**A**), green light (**B**), red light (**C**) and far-red light (**D**) under 25 and 100 µmol photons $m^{-2} \cdot s^{-1}$. Values represent means of four biological independent replicates and asterisks show statistical difference compared to wild type according to Student's *t* test (two-sided, $P < 0.05$)**.**

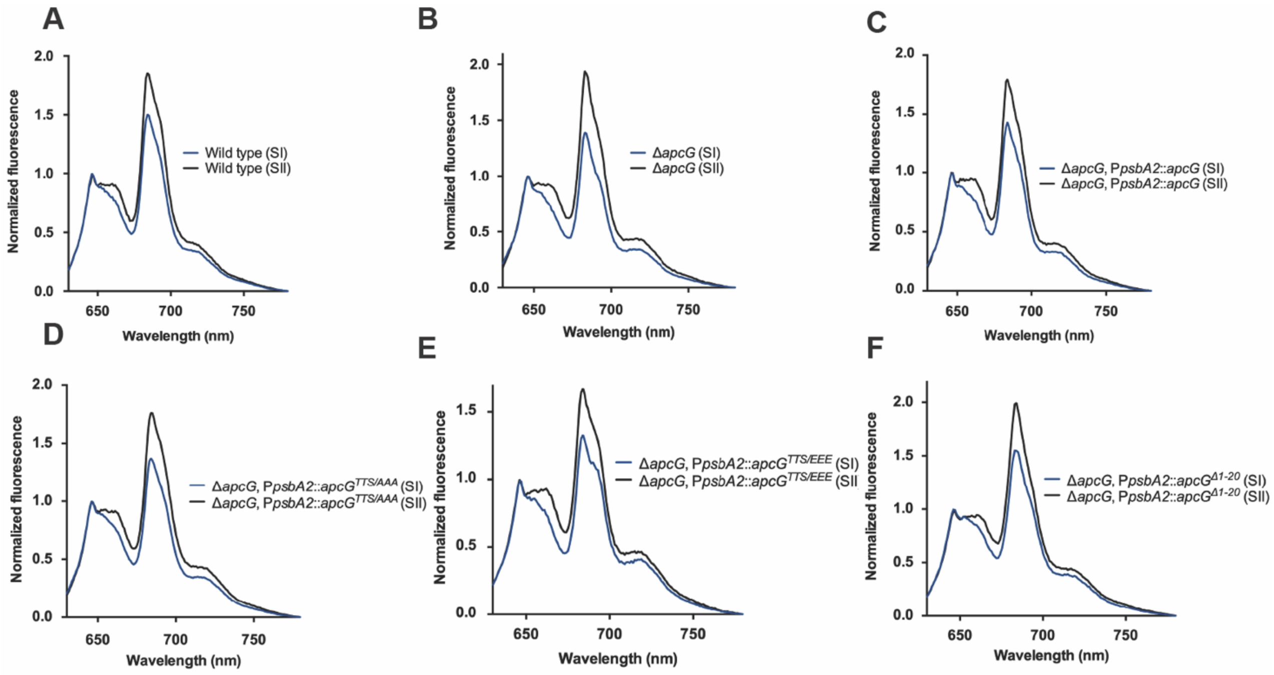


**Supplemental figure S4. ApcG does not play a role in state transitions.** Low temperature (77 K) fluorescence spectra from *Synechocystis* cultures were obtained under state I (induced with blue light pre-treatment) and state II (pre-treated under darkness). The strains used correspond to *Synechocystis* wild type (**A**), *apcG* deletion (**B**), *apcG* wild type complementation (**C**), permanent non-phosphorylated *apcG*$^{TTS/AAA}$ (**D**), phospho-mimicking *apcG*$^{TTS/EEE}$ (**E**) and truncated *apcG*$^{\Delta 1\text{-}20}$ (**F**). Values correspond to the mean of three biological replicates. Spectra were normalized to the peak of PBS at 646 nm.

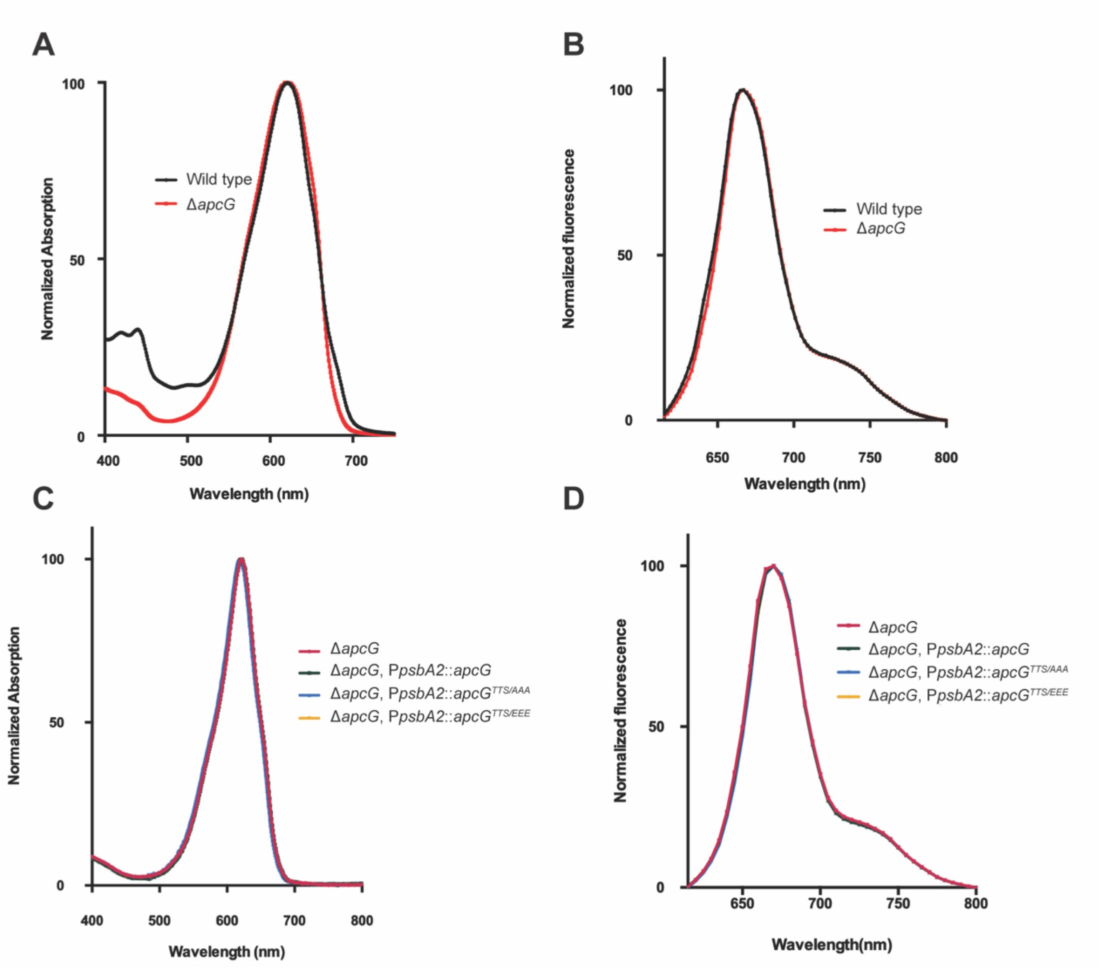


**Supplemental figure S5. Absorption and fluorescence spectra of isolated PBS from deletion and complementation strains.** PBS isolated after sucrose gradient were used to record their absorption and fluorescence spectra to assess their integrity. The wild type and *apcG* deletion strains absorption and fluorescence spectra are shown in **(A)** and **(B)** respectively. Additionally, comparison of PBS obtained from the deletion and complementation strains are shown in their respective absorption **(C)** and fluorescence **(D)** spectra. Values correspond to technical triplicates (normalized to their maxima) from a representative experiment out of three biologically independent samples.

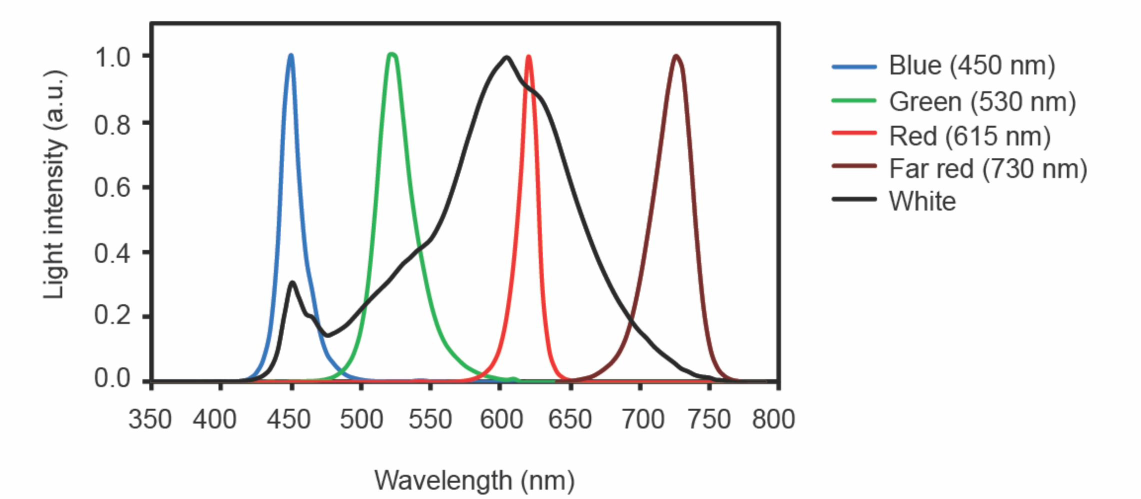


**Supplemental figure S6. Light sources spectra.** The light sources spectra used for growing cyanobacteria strains were recorded from 350 to 800 nm. Light intensity is represented as arbitrary units (a.u.) normalized to each color maxima.
